# A novel effective live-attenuated human metapneumovirus vaccine candidate produced in the serum-free suspension DuckCelt®-T17 cell platform

**DOI:** 10.1101/2021.08.30.458186

**Authors:** Caroline Chupin, Andrés Pizzorno, Aurélien Traversier, Pauline Brun, Daniela Ogonczyk-Makowska, Blandine Padey, Cédrine Milesi, Victoria Dulière, Emilie Laurent, Thomas Julien, Marie Galloux, Bruno Lina, Jean-François Eléouët, Karen Moreau, Marie-Eve Hamelin, Olivier Terrier, Guy Boivin, Julia Dubois, Manuel Rosa-Calatrava

**Author notes:** These authors contributed equally (Caroline Chupin, Andrés Pizzorno). These authors contributed equally (Julia Dubois, Manuel Rosa-Calatrava).

## Abstract

Human metapneumovirus (HMPV) is a major pediatric respiratory pathogen for which there is currently no specific treatment or licensed vaccine. Different strategies have been evaluated to prevent this infection, including the use of live-attenuated vaccines (LAVs). However, further development of LAV approaches is often hampered by the lack of highly efficient and scalable cell-based production systems that support worldwide vaccine production. In this context, avian cell lines cultivated in suspension are currently competing with traditional cell platforms used for viral vaccine manufacturing. We investigated whether the DuckCelt®-T17 avian cell line (Vaxxel) we previously described as an efficient production system for several influenza strains could also be used to produce a new HMPV LAV candidate (Metavac®), an engineered SH gene-deleted mutant of the A1/C-85473 strain of HMPV. To that end, we characterized the operational parameters of multiplicity of infection (MOI), cell density, and trypsin addition to achieve optimal production of the LAV Metavac® in the DuckCelt®-T17 cell line platform. We demonstrated that the DuckCelt®-T17 cell line is permissive and is well adapted to the production of the wild-type A1/C-85473 HMPV and the Metavac® vaccine candidate. Moreover, our results confirmed that the LAV candidate produced in DuckCelt®-T17 cells conserves its advantageous replication properties in LLC-MK2 and 3D-reconstituted human airway epithelium models, as well as its capacity to induce efficient neutralizing antibodies in a mouse model. Our results suggest that the DuckCelt®-T17 avian cell line is a very promising platform for scalable in-suspension serum-free production of the HMPV-based LAV candidate Metavac®.

## Introduction

Human pneumoviruses, which include the human respiratory syncytial virus (HRSV) and the human metapneumovirus (HMPV), are a major worldwide cause of acute respiratory tract infections, especially among children, older adults, and immunocompromised individuals ^1–3^. Infections by these two respiratory pathogens share many features and are globally responsible for more than 33 million annual cases among children under 5 years old ^3–5^. Despite this important clinical burden, there is currently no licensed vaccine or specific antiviral against human pneumoviruses. To date, only one humanized monoclonal antibody (Palivizumab) was regulatory approved for the passive immunoprophylaxis against severe HRSV infection in high-risk infants and children ^6,7^.

Throughout the last decades, several vaccine strategies have been developed in order to prevent disease caused by pneumoviruses, mostly based on recombinant proteins, live-attenuated vaccines (LAVs) ^8,9^ or, more recently, mRNA candidates ^10–12^. Of note, the LAV strategies are considered to be well adapted to pediatric immunization and have the advantage of eliciting both humoral and mucosal immunity by mimicking natural viral replication routes, in addition to being delivered without adjuvant ^13^. This contrasts with formalin-inactivated pneumovirus-based vaccines, which in the past have been responsible for events of vaccine-induced enhanced disease ^14^. Some pneumovirus-based LAV candidates led to promising outcomes in *in vitro* and *in vivo* experiments, such as M2-2, NS2, and G/SH gene-deleted HRSVs, and G/SH-deleted HMPVs ^9,15,16^. Unfortunately, final reports on some of these candidates show them to be over-attenuated and/or ineffective at inducing protective antibody response in human clinical trials ^8,9,17,18^. Only a small number of HMPV LAV candidates have shown the potential to progress towards clinical evaluation ^8,15,19,20^. In this context, we have previously described an engineered SH gene-deleted recombinant virus based on the hyperfusogenic A1/C-85473 HMPV strain (ΔSH-rC-85473) as a promising LAV candidate (Metavac®): it has shown efficient replication in a human cell-based system, as well as protective properties in mice lethally challenged with wild type HMPV, including the induction of neutralizing antibodies, reduced disease severity, weaker inflammatory responses, and a balanced stimulation of the immune response ^20^.

On the other hand, development of LAV-based strategies is often hampered by the limited availability of highly efficient and scalable cell-based production platforms to support the vaccine need. Currently, viral vaccine production is mostly performed using adherent cells, such as the Vero and MRC5 cell lines ^21–23^. These cell lines are well known and grown in roller bottles or multiplate cell factory systems ^22^. However, costs, space and workforce constraints prevent these technologies from being easily scalable. In this context, serum-free suspension cell lines such as the human cell lines PER.C6 (Crucell) ^24^ and CAP® (CEVEC Pharmaceutical) ^25^ or the avian cell lines AGE1.CR® (Probiogen) ^26^, EB66® (Valneva) ^27^, and QOR/2E11 cells (Baxter) have been developed to ease cell culture and viral amplification steps ^21^. These cell lines enable reduction of footprint and labor intensiveness, as they are cultured in bioreactors without microcarriers. During evaluation into whether these have the potential to become versatile viral production platforms, the AGE1.CR® and EB66® avian cell lines have been shown to efficiently produce several viruses, such as Modified Vaccinia Ankara MVA or influenza viruses ^22,26–29.^ In contrast, none of these new-generation cell lines has yet been reported to be permissive or to support production of pneumoviruses. Hence, HRSV is commonly cultured onto anchorage-dependent human Hep-2 cells ^30^ and HMPV onto adherent non-human primate LLC-MK2 cells ^31^.

Considering scalability in the manufacturing process of the *Cairina moschata* duck embryo-derived DuckCelt®-T17 cell line (Vaxxel), which we previously described as an efficient platform for production of human and avian influenza viruses ^28^, we sought to evaluate its putative permissiveness and capacity to produce C-85473 HMPV-based viruses, notably our new LAV candidate Metavac® ^20^. We characterized the main operational parameters for viral production, including multiplicity of infection (MOI), cell density, and trypsin input to achieve optimal production yield. Finally, using *in vitro* and *in vivo* experimental models, we highlighted the conservation of morphological features, replicative capacities, and immunizing properties of the Metavac® virus produced in the in-suspension DuckCelt®-T17 cell line.

## Results

### The DuckCelt®-T17 cell line is permissive to C-85473 HMPV and is appropriate for viral production

Firstly, we sought to evaluate the permissiveness of DuckCelt®-T17 cells to the prototype WT C-85473 HMPV and its recombinant GFP-expressing reporter counterpart (rC-85473-GFP) in routine cell culture parameters, as previously described ^28^. As HMPV F protein cleavage and related virus propagation are known to be trypsin-dependent in adherent cell culture ^32^, we supplemented the culture medium with acetylated trypsin at the final concentration of 0.5 μg/mL at the time of infection (D0) and two, four and seven days post-infection (D2, D4, and D7, respectively).

Thanks to the follow-up of reporter GFP gene expression in the cells infected by the recombinant HMPV (rC-85473-GFP), we observed efficient virus propagation in the cellular suspension over a 10-day period, hence validating the permissiveness of DuckCelt®-T17 cells to HMPV infection and replication. At a MOI of 0.01, we observed by fluorescent microscopy maximal GFP expression at 7 days post-infection (7 dpi), as illustrated in **Figure 1a**. Viral kinetics of WT C-85473 and rC-85473-GFP viruses were also characterized by measuring virus production from culture supernatant. We measured mean maximum viral titers of 9.8×10^6^ TCID_50_/mL at 6 dpi with the WT C-85473 strain and 1.9×10^6^ TCID_50_/mL at 8 dpi with the rC-85473-GFP virus (**Figure 1b**). Accordingly, DuckCelt®-T17 cells achieved a maximal cell density of 5.5×10^6^ cell/mL at 4 dpi or 6.6×10^6^ cell/mL at 6 dpi when infected with WT C-85473 or rC-85473-GFP viruses, respectively (**Figure 1c**). In comparison, mock-infected cell suspension achieved a maximal cell concentration of 8.6×10^6^ cell/mL after 8 days of culture (**Figure 1c).**Thus, these results indicate that the DuckCelt®-T17 cell line is permissive to C-85473 HMPV-based viruses and allows their efficient amplification by 2 log_10_ compared to the initial inoculum within an 8-day period.

**Figure 1.**
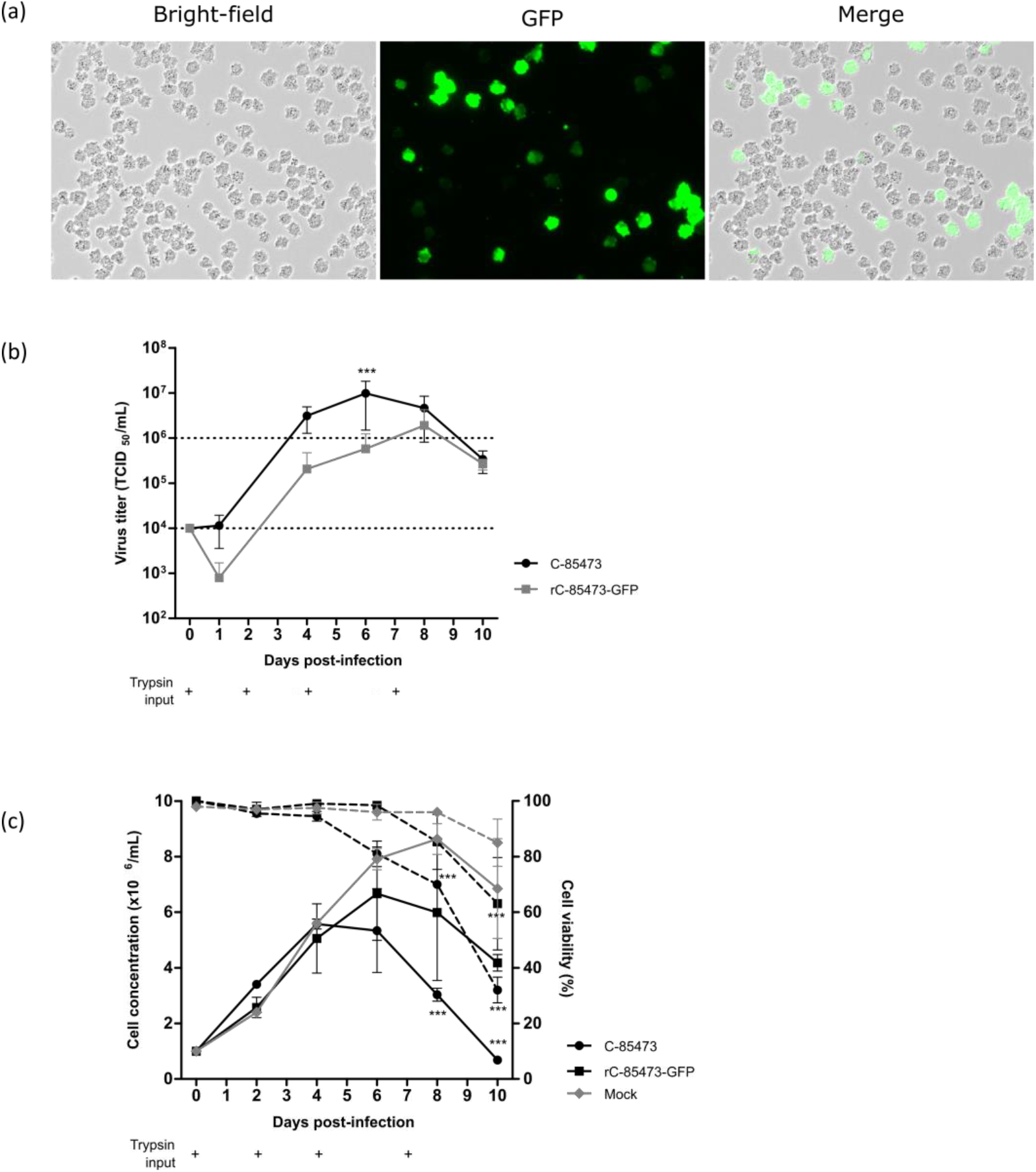
Viral kinetics of the wild-type C-85473 HMPV and recombinant rC-85473-GFP HPMV in the DuckCelt®-T17 cell line. Cells were seeded at 1×10^6^ cell/mL in a 10mL working volume and infected at a MOI of 0.01. Trypsin was added at 0.5 μg/mL on Day (D) 0, D2, D4, and D7 (+). (**a**) Picture of T17 cells infected with rC-85473-GFP at 7 dpi (days post-infection). Each picture was taken in bright field and fluorescent microscopy (x20 magnification). (**b**) Viral titers were measured from culture medium as 50% tissue culture infectious doses (TCID_50_)/mL in LLC-MK2 cells. (**c**) Cell growth (solid line) and viability percentage (dotted line) were measured with the Countess™ II FL Automated Cell Counter. Results are shown as means ± SD and represent duplicates in two independent experiments. *** p < 0.001 when comparing the infected conditions to each other (**b**) or to the mock condition (**c**) using a two-way repeated measures ANOVA.

### Identification of best operating parameters for viral production of the rC-85473-GFP virus in the DuckCelt®-T17 cell line

Starting from the standard viral culture parameters mentioned above, we aimed to evaluate separately the influence of the MOI, cell density, time, and concentration of trypsin input on the HMPV yield in order to identify key operating parameters for the production process in a 10 mL working volume of DuckCelt®-T17 cells.

First, DuckCelt®-T17 cells were seeded at 1×10^6^ cell/mL in supplemented OptiPRO™ SFM and inoculated the same day with rC-85473-GFP HMPV at three different MOI (0.1, 0.01, and 0.001). Whereas at a MOI of 0.1, viral production flattened between 6 and 10 dpi with mean virus titers of 2.4×10^6^ TCID_50_/mL, mean virus titers were between 2.08 and 6.58×10^6^ TCID_50_/mL for infection at a MOI of 0.01 at the same time point (**Figure 2a**). In comparison, when an even lower MOI of 0.001 was used, we measured a maximum virus yield of 0.78×10^6^ TCID_50_/mL after 10 dpi, significantly lower and later than with a MOI of 0.01 and 0.1 (**Figure 2a**). Moreover, percentage cell infectivity, determined by quantification of GFP-positive cells by flow cytometry, showed that nearly 100% of DuckCelt®-T17 cells were infected at 6 dpi at a MOI of 0.1, 8 dpi at a MOI of 0.01, and 10 dpi at a MOI of 0.001 (**Figure 2b**). Hence, despite faster viral replication kinetics in the cellular suspension, the use of a tenfold higher MOI did not significantly increase virus production compared to MOI 0.01.

**Figure 2.**
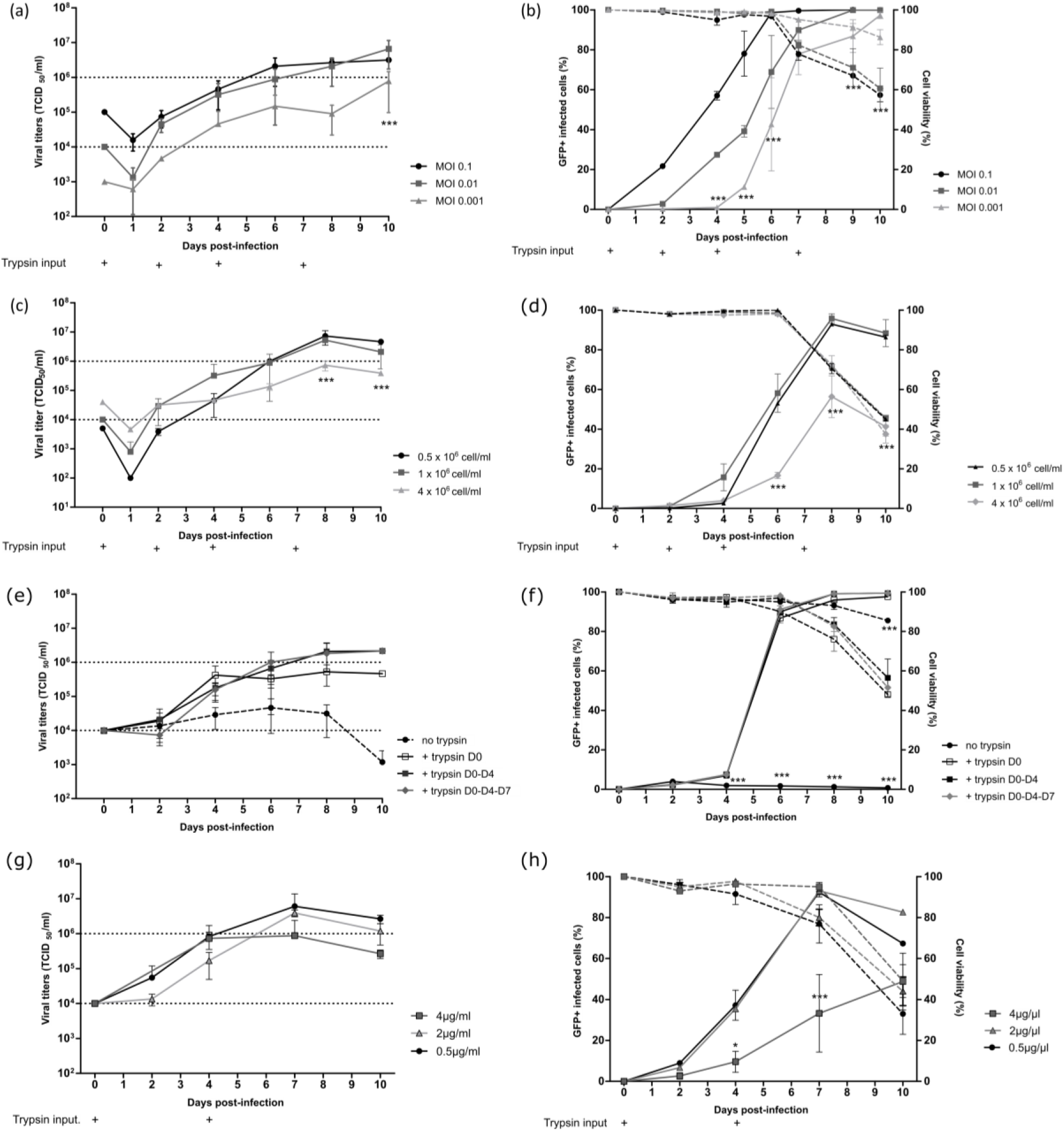
Evaluation of multiplicity of infection (MOI), cell density, trypsin concentration, and optimal timing for trypsin addition on rC-85473-GFP HMPV production kinetics in the DuckCelt®-T17 cell line. Culture parameters in a 10 mL working volume were evaluated separately by viral titration (**a–c–e–g**) and cell viability and infectivity measurement (**b–d–f–h**; dotted line for viability and solid line for infectivity). Viral titers were measured from culture medium as TCID50/mL in LLC-MK2 cells. Viability was measured with trypan blue using an automated cell counter. Infected GFP-positive cells were evaluated with the FACS CantoII. (**a–b**) Evaluation of a MOI of 0.1, 0.01 or 0.001. (**c– d**) Evaluation of three different cell densities at the time of infection: 0.5, 1, or 4×10^6^ cell/mL (**e–f**) Evaluation of timing of trypsin addition. Trypsin was added at 0.5 μg/mL at D0, at D0 and D4, or at D0, D4, and D7, or no trypsin was added. (**g–h**) Evaluation of trypsin concentration: 0.5, 2, or 4 μg/mL. Results are shown as means ± SD and represent duplicates in two independent experiments. * p < 0.05, ** p < 0.01, *** p < 0.001 when comparing infected conditions to each other using a two-way repeated measures ANOVA.

We then considered the influence of the cell density at the time of virus inoculation. DuckCelt®-T17 cells were centrifuged in order to be seeded at three different cell concentrations: 0.5, 1, and 4×10^6^ cells/mL in OptiPRO™ SFM (50% conditioned medium and 50% fresh medium). Cell suspensions were then inoculated with rC-85473-GFP HMPV at a MOI of 0.01 and supplemented with acetylated trypsin at 0.5 μg/mL. Maximum virus yields of 5.26×10^6^ and 7.32×10^6^ TCID_50_/mL were achieved at 8 dpi when 0.5 or 1×10^6^ cells/mL, respectively, were inoculated (**Figure 2c**). In contrast, significantly lower mean peak viral titers (0.73×10^6^ TCID_50_/mL) were measured at 8 dpi when using an initial cell concentration of 4×10^6^ cells/mL (**Figure 2c**), in line with the observed significant reduction in the maximal percentage of GFP-positive cells. These results show that increasing cell density above 1×10^6^ cells/mL at the time of inoculation results in no benefit for HMPV production (**Figure 2d**).

Finally, we looked at the impact of trypsin on virus yield by testing repeated supplementation or increasing its concentration in the cell culture medium. DuckCelt®-T17 cells were seeded at 1×10^6^ cell/mL and then inoculated with rC-85473-GFP HMPV at MOI 0.01. The culture medium was supplemented or not at varying time points (D0, D0 and D4, or D0, D4, and D7) with 0.5 μg/mL acetylated trypsin (**Figure 2e-f**). When comparing cell infectivity and virus titers between experimental conditions, we confirmed that trypsin supplementation is necessary for viral replication in DuckCelt®-T17 cells, as illustrated by the absence of both virus amplification and GFP-positive cells in the cell suspension in the absence of trypsin (**Figure 2e-f**). Efficient and comparable virus propagation was observed after the addition of trypsin at one, two, or three time points, resulting in nearly 100% of cells being infected at 8 dpi (**Figure 2f**). However, viral production and release in the culture medium seemed to be impaired when trypsin was only added on D0, as reflected by overall viral yields that were at least tenfold lower between 8 and 10 dpi in comparison with conditions when trypsin was also supplemented at D4 (**Figure 2e**). Interestingly, a third addition of trypsin at D7 did not increase virus production (**Figure 2e**).

Based on these results, we further evaluated supplementation of the culture medium with increasing concentrations of trypsin, notably 0.5, 2 or 4 μg/mL (**Figure 2g-h**). In accordance with the low percentage of infected cells detected (**Figure 2h**), while the addition of 4 μg/mL of acetylated trypsin did not result in virus titers higher than 1×10^6^ TCID_50_/mL, the addition of 0.5 or 2 μg/mL trypsin led respectively to a 6.5-fold or 4.5-fold higher peak of viral production at 7 dpi (**Figure 2g**).

In conclusion, we identified the best operating parameters to amplify the rC-85473-GFP HMPV in a 10 mL working volume of DuckCelt®-T17 cells (1×10^6^ cells/mL at the time of inoculation with a MOI of 0.01 and two additions of 0.5μg/mL acetylated trypsin, on D0 and D4), leading to 2 log_10_ higher production yield in comparison to the initial inoculum.

### Production of the Metavac® LAV candidate in the DuckCelt®-T17 cell line using optimized operating parameters

We further aimed to determine if the best operating production parameters identified with the rC-85473-GFP virus in DuckCelt®-T17 cells were well adapted to the production of our previously described novel LAV candidate Metavac® ^20^, which is an engineered SH gene-deleted version of the C-85473 strain of HMPV. We therefore inoculated 1×10^6^ DuckCelt®-T17 cells/mL with Metavac® at MOIs of 0.1, 0.01, or 0.001 in cell culture medium supplemented with 0.5μg/mL of acetylated trypsin on D0 and D4 post-infection. As shown in **Figure 3a**, the maximum viral production was achieved at 10 dpi with 1.02×10^6^ TCID_50_/mL, 1.94×10^6^ TCID_50_/mL, or 0.62×10^6^ TCID_50_/mL for a MOI of 0.1, 0.01, or 0.001, respectively. When we measured the percentage of infected cells, the maximum infectivity was achieved in 6 days at a MOI of 0.1, similar to timings observed with the rC-85473-GFP virus (**Figure 2b**), whereas 10 days were necessary to infect the whole cell suspension at a MOI of 0.01 (**Figure 3b**). Interestingly, a MOI of 0.001 was not sufficient to allow propagation of the Metavac® virus to more than 36.2 ± 37.7% of cells after 10 days of culture.

**Figure 3.**
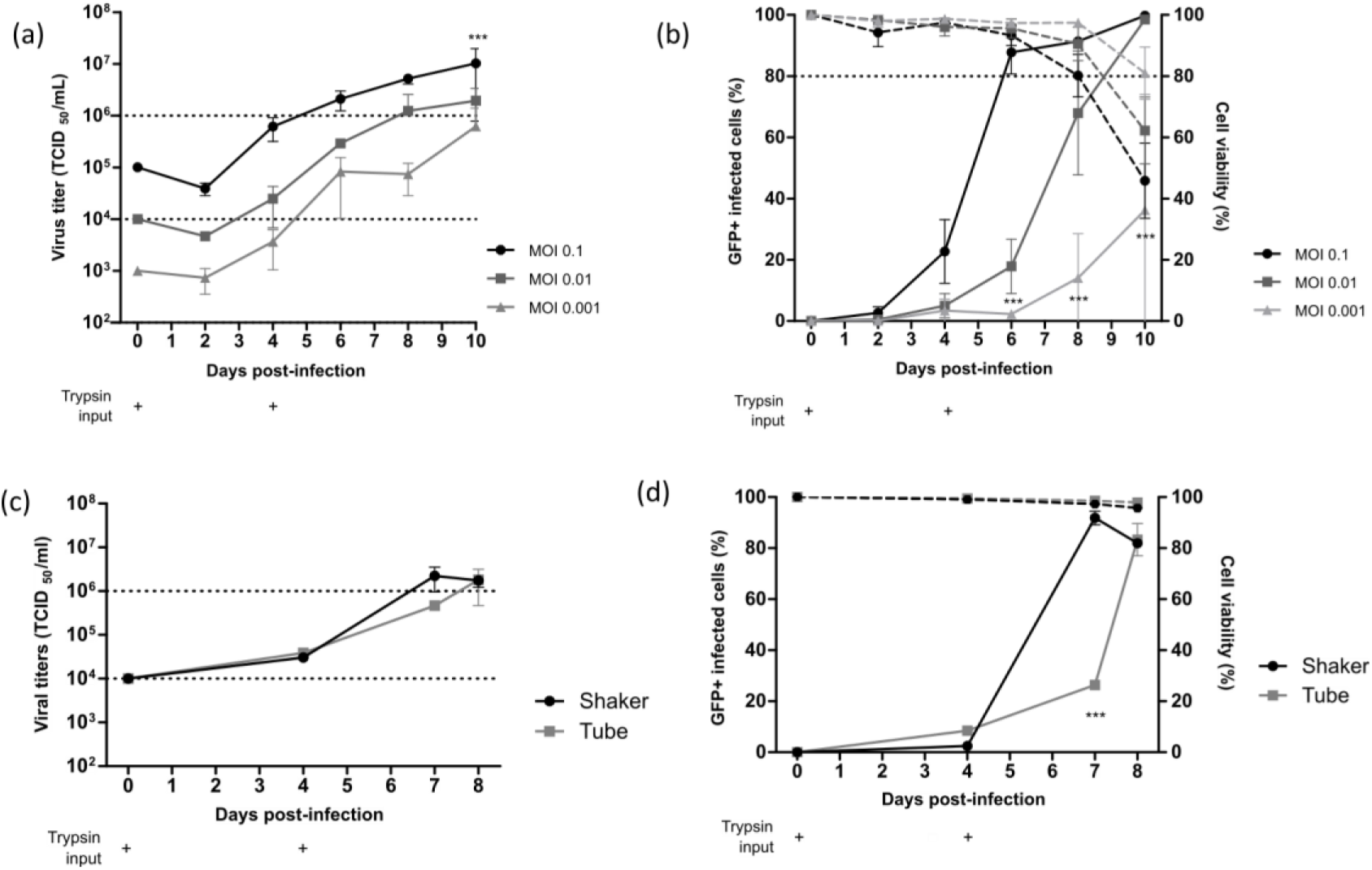
Viral kinetics of live-attenuated vaccine candidate ΔSH-rC-85473-GFP HMPV in the DuckCelt®-T17 cell line. Based on optimal production parameters identified, cells were seeded at 1×10^6^ cell/mL, infected, and trypsin was added at 0.5μg/mL on D0 and D4 (**+**).Culture parameters in a 10 mL working volume were evaluated separately by viral titration (**a–c**) and viability and infectivity measurement (**b–d**; dotted line for cell viability and solid line for infectivity). **(a–b)** Evaluation of viral kinetics and cell infectivity for a MOI of 0.1, 0.01, or 0.001 in a 10 mL working volume. (**c–d**) Evaluation of viral kinetics and cell infectivity when cells were seeded in a 500 mL (shaker) or 10 mL (tube) working volume and infected at a MOI of 0.01. Results are shown as means ± SD and represent duplicates in two independent experiments. *** p < 0.001 when comparing infected conditions to each other using a two-way repeated measures ANOVA.

To investigate the scalability of the DuckCelt®-T17 production platform, we then performed Metavac® production in a 500 mL working volume in shaker flasks, using the selected best operating parameters and a MOI of 0.01. Similar to the results obtained in 10 mL cultures, peak virus production was 2.2×10^6^ TCID_50_/mL at 7 dpi (**Figure 3c**), which corresponded with the maximum percentage of infected GFP-positive cells (**Figure 3d)**.

Altogether, these results show that the DuckCelt®-T17 cell line is permissive and well adapted to scalable production of our LAV candidate. However, as Metavac® is an attenuated virus, the duration of the production process could be longer than with the rC-85473-GFP strain but is expected to reach comparable production yields.

### HMPV virions produced in the DuckCelt®-T17 cell line conserve their morphological characteristics and full replicative properties in LLC-MK2 cells and reconstituted HAE

We further characterized key morphological and functional viral properties of rC-85473-GFP and Metavac® viruses produced in DuckCelt®-T17 cells. Transmission electron microscopy analysis of virions released in the culture medium revealed typical HMPV pleiomorphic virus particles with a mean diameter of 174.4 nm and 183.2 nm for rC-85473-GFP and Metavac®, respectively, presenting abundant glycoproteins at their surface (**Figure 4a**). Viruses produced in the DuckCelt®-T17 cell line were then assessed for their replicative properties in LLC-MK2 cells over a 7-day period (**Figure 4b-d**). In line with previous studies using the viral hyperfusogenic C-85473 background ^33,34^, fluorescence microscopy showed the formation of large syncytia, clearly visible as early as 4 dpi (**Figure 4b**). Peak viral titers of approximately 4.95×10^5^ TCID_50_/mL and 4.15×10^5^ TCID_50_/mL were reached by 5 dpi for rC-85473-GFP and Metavac® viruses, respectively (**Figure 4c**). Additionally, crystal violet coloration of infected LLC-MK2 monolayers revealed very similar kinetics between both DuckCelt®-T17-produced rC-85473-GFP and Metavac® viruses and those produced in LLC-MK2 (**Figure 4d**).

**Figure 4:**
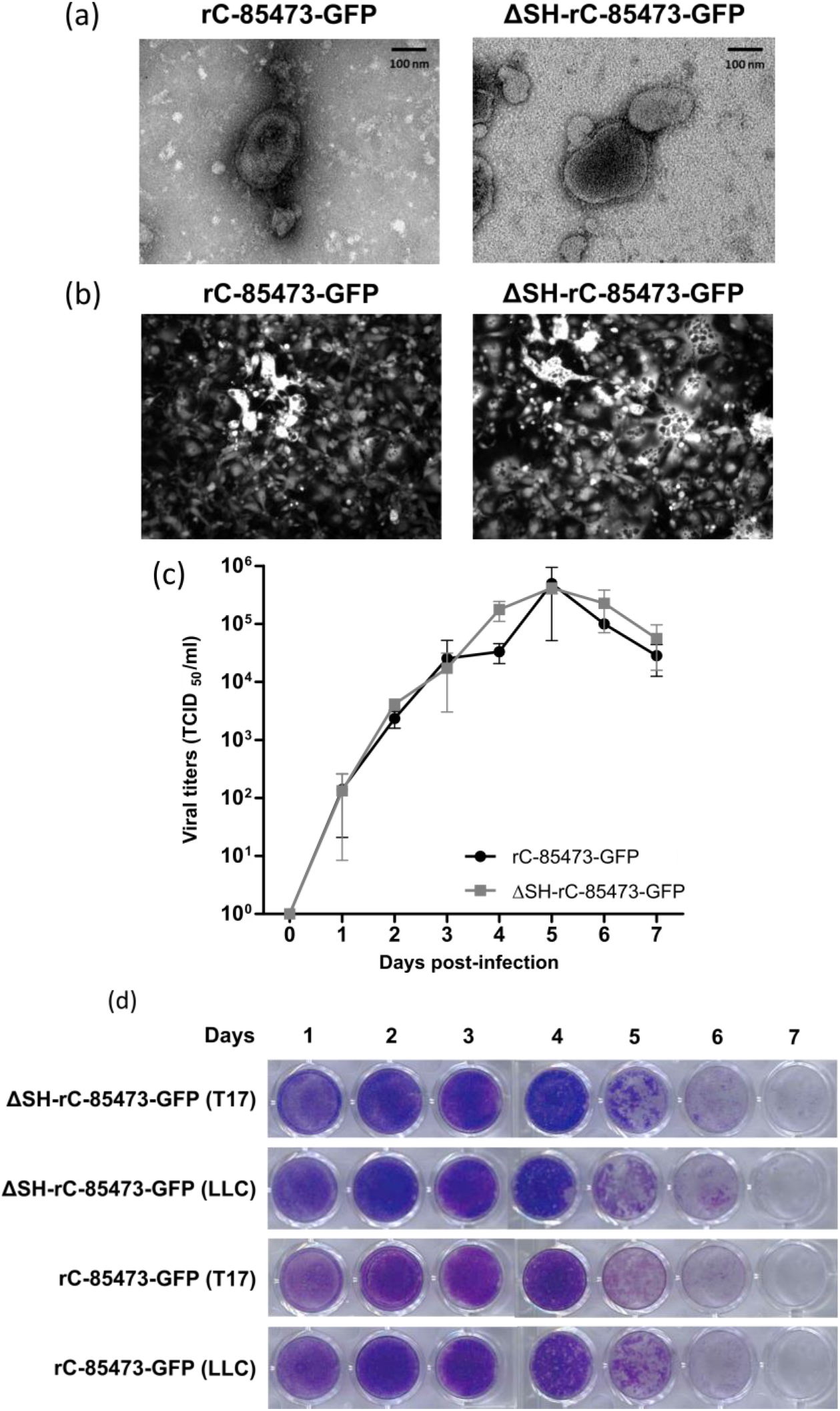
Visualization of T17-produced rC-85473-GFP and ΔSH-rC-85473-GFPrHMPV virus particles and evaluation of *in vitro* replicative capacity. **(a)** Representative negative stain electron microscopy images of rC-85473-GFP and ΔSH-rC-85473-GFP virions, obtained from DuckCelt®-T17 culture, are presented. Bar represents 100 nm. **(b-d)** LLC-MK2 monolayers in 24-well plates were infected with each of the recombinant HMPVs at a MOI of 0.01. **(b)** Images of representative cytopathic effects of each virus were captured after 4 dpi by fluorescent microscopy (x10 magnification). **(c)** Supernatants were harvested every 24 h for 7 days, frozen and subsequently thawed and titrated as TCID_50_/mL onto LLC-MK2 cells. Growth curves represent mean titers ± SD of each time point titrated in triplicate. **(d)** Infected LLC-MK2 monolayers were fixed in formaldehyde after harvest and images were captured after crystal violet coloration.

We then evaluated rC-85473-GFP and Metavac® viruses using the Mucilair™ 3D-reconstituted HAE *in vitro* physiological model of infection ^20,35^. Both viruses were able to infect and fully spread within the HAE, as shown by their reporter GFP expression pattern at 5 dpi (**Figure 5a** and **5b**). Moreover, we measured the progeny virus secretion at the HAE apical surface during the time course of infection by quantification of N gene copies (**Figure 5c**). In line with previous results ^20^, both DuckCelt®-T17-produced viruses demonstrated high replicative capacity in HAE. Viral amplification occurred mainly between 2 dpi and 5 dpi, but viral persistence was observed until at least 12 dpi. Maximal apical viral titers of 1.07×10^9^ and 1.29×10^9^ N copies per apical sample were reached at 5 dpi for rC-85473-GFP and Metavac®, respectively (**Figure 5c**).

**Figure 5:**
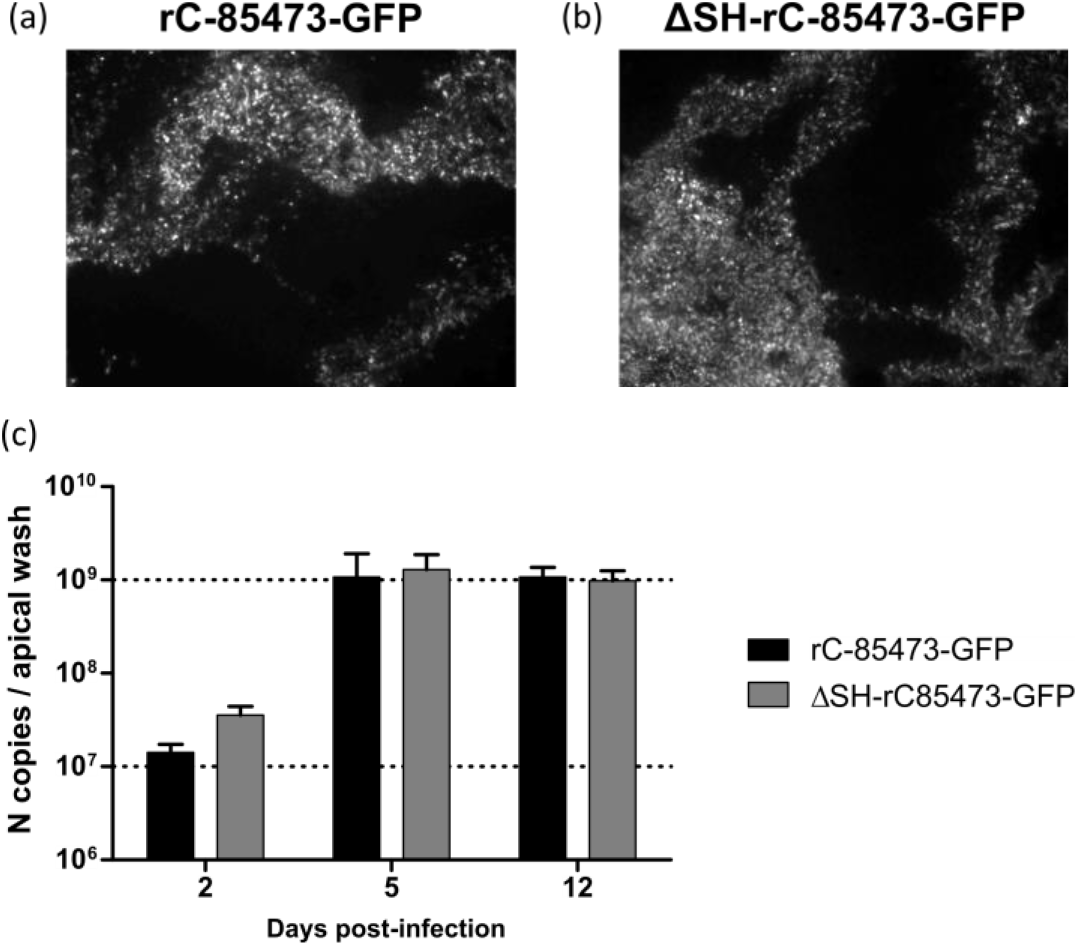
Recombinant HMPVs produced in the DuckCelt®-T17 cell line conserved infectivity and replicative capacity in 3D-reconstituted human airway epithelium (HAE). MucilAir™ epithelium from healthy donors were infected with rC-85473-GFP or ΔSH-C-85473-GFP viruses produced in DuckCelt®-T17 cells at a MOI of 0.1 and monitored for 12 days. Viral spread of rC-85473-GFP T17 (**a**) or ΔSH-C-85473-GFP T17 (**b**) in HAE at 5 dpi was observed by fluorescence microscopy (10x magnification). Viral quantification from epithelium apical washes (**c**) at 2, 5, and 12 dpi was performed by specific RT-qPCR of the N viral gene. Data are shown as means ± SD and represent experimental duplicates.

Taken together, our results indicate that rC-85473-derived HMPVs produced in the in-suspension avian DuckCelt®-T17 cell platform fully conserve their *in vitro* phenotype and harbor efficient viral replication in both LLC-MK2 and HAE models.

### Metavac® vaccine candidate produced in DuckCelt®-T17 cells conserves full immunogenic properties in mice

Considering the potential of the Metavac® virus as a HMPV LAV candidate, we further evaluated its capacity to infect, replicate in, and immunize BALB/c mice. We therefore infected BALB/c mice intranasally with 1×10^6^ TCID_50_ of Metavac® either produced in LLC-MK2 or DuckCelt®-T17 cells. In agreement with previous results ^20,36^, neither weight loss nor clinical signs were observed during the 14-day follow-up in the two Metavac®-infected groups, compared to the non-infected (mock) group (**Figure 6b**). After 5 and 14 dpi, we quantified the viral pulmonary replication by RT-qPCR from lung homogenates (**Figure 6a**). Metavac® viruses produced in either LLC-MK2 or DuckCelt®-T17 cells replicated efficiently in the lungs of infected mice after 5 dpi and were almost cleared by 14 dpi (**Figure 6c**), as previously described for LAVs based on a C-85473 strain of HMPV in which the SH gene is deleted ^20,36^.

**Figure 6:**
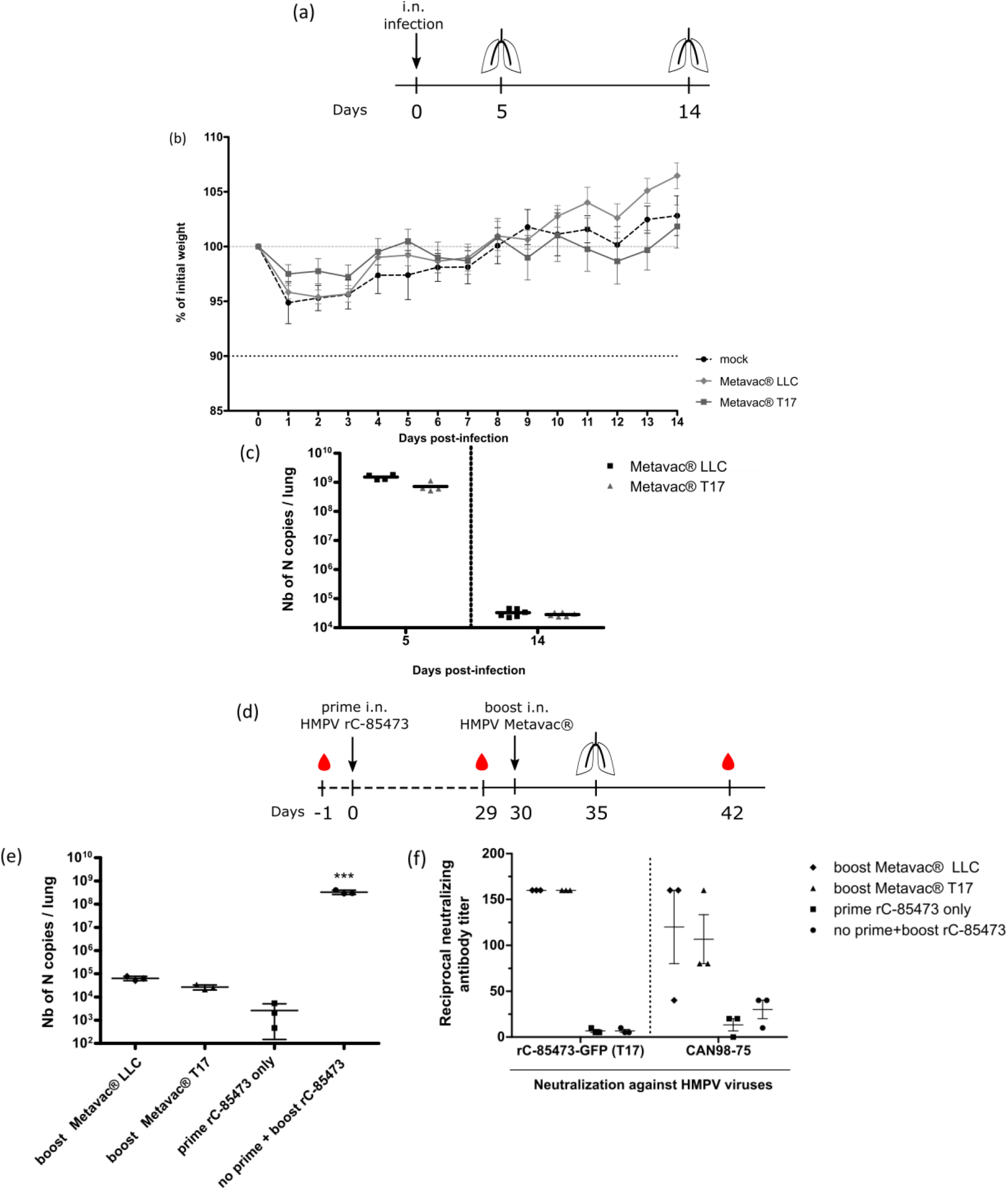
Evaluation of pulmonary replication and immunogenicity in BALB/c mice after prime or boost infection with the T17-produced HMPV LAV candidate. (**a–c**) BALB/c mice were intranasally infected with 1×10^6^ TCID_50_ of the ΔSH-rC-85473-GFP virus and monitored for 14 days after infection (n = 20), as represented in schematic timeline (**a**), with weight loss (**b**, n=16) and pulmonary viral titers quantified by RT-qPCR from lungs harvested at 5 dpi (**c**, n = 4). (**d-f**) BALB/c mice were prime-infected with 5×10^5^ TCID_50_ of the rC-85473-GFP virus, then boost-infected after 30 dpi with 5×10^5^ TCID_50_ of the ΔSH-rC-85473-GFP LLC or T17 viruses *via* the intranasal route and monitoring was performed as presented in (**d**). (**e**) Pulmonary viral titers quantified by RT-qPCR from lungs harvested at 5 days post-boost (n = 3). (**f**) Induction of neutralizing antibodies by Metavac® viruses in mice 21 days post-boost. Three pools of sera from two mice were tested for neutralization against two WT HMPV strains, the homologous rC-85473-GFP virus produced in DuckCelt®-T17 cells and the heterologous CAN98-75 virus, resulting in three biological replicates per group. One day before prime infection, the naïve status of mice was confirmed by a microneutralization assay from a pool of sera. *** p < 0.001 when comparing each group to the no-boost condition using a one-way ANOVA with Dunnett’s post-test. i.n.: intranasal.

We finally investigated the capacity of Metavac® to induce the production of high levels of HMPV neutralizing antibodies *in vivo* (**Figure 6d**). BALB/c mice were prime-infected with 5×10^5^ TCID_50_ of rC-85473-GFP and boost-infected 30 days later *via* the intranasal route with 5×10^5^ TCID_50_ of Metavac® viruses either produced in LLC-MK2 or DuckCelt®-T17 cells. Blood samples were collected prior to prime and boost, as well as 21 days post-boost (**Figure 6d)**. As expected, we detected low levels of viral genome in the lungs of all groups that received prime and boost intranasal infections (1×10^4^ to 1×10^5^ of N gene copies per lung, see **Figure 6e**), in contrast with the group prime-instilled with OptiMEM medium and boost-infected with rC-85473-GFP (up to 5×10^8^ of N gene copies per lung, **Figure 6e**).

In line with these results, high levels of neutralizing antibodies were titrated 21 days after boost by both Metavac® viruses, in comparison with control groups that received either prime or boost rC-85473-GFP infections (**Figure 6f**). Interestingly, three weeks after boost, sera from both groups could efficiently neutralize homologous rC-85473-GFP HMPVs produced in DuckCelt®-T17 cells, and the heterologous CAN98-75 strain, whereas no significant HMPV antibody response was measured 29 days after only the prime infection in all groups of mice, as expected. In addition, rC-85473-GFP HMPVs produced in LLC-MK2 cells were similarly neutralized by antibodies induced by both boosts with Metavac® viruses produced in LLC-MK2 or DuckCelt®-T17 cells (**Supplementary Table 1**).

Altogether, our results indicate that the Metavac® vaccine candidate produced in the in-suspension avian DuckCelt®-T17 cell platform conserves full immunization properties by inducing efficiently neutralizing antibodies against both homologous and heterologous WT patient-derived strains in a murine model.

## Discussion

Despite the worldwide burden of human pneumoviruses and the effort to develop vaccine strategies, there is still no approved vaccine against HRSV or HMPV. When considering the production of LAV candidates, one of the major obstacles is the deficit of scalable cell lines able to respond to industrial production requirements. In this study, we confirmed that the DuckCelt®-T17 cell line, which we have already described for its serum-free suspension cultivation and its ability to efficiently produce influenza viruses ^28^, can respond to the need for a scalable cell line for the manufacture of HMPV, and more specifically of our LAV candidate Metavac®, derived from the C-85473 strain of HMPV ^20^. We demonstrated that the DuckCelt®-T17 cell line supports Metavac® replication with high yields in upscalable cultivation conditions, conserving both *in vitro* replication properties (in LLC-MK2 and 3D-reconstituted HAE models) and the ability to infect and induce a neutralizing antibody response in a mouse model.

Since the first description of HMPV in 2001, a limited number of cell-based production platforms, such as Vero or LLC-MK2 cells, have been shown to support HMPV replication. Moreover, in adherent cell lines, pneumoviruses are well known to spread preferentially by different cell-to-cell mechanisms ^37^. Given these considerations, it is of particular interest that the DuckCelt®-T17 cell platform is susceptible to C-85473 HMPV infection and efficiently supports its propagation into the cellular suspension, in culture conditions compatible with the in-suspension properties of the DuckCelt®-T17 cell line. Of note, the virus particles are directly detected and titrated from the culture medium without requiring mechanical cell lysis and exhibit the expected morphological features. Moreover, in this study, we have shown that the vaccine candidate Metavac® produced in DuckCelt®-T17 cells conserved its replicative properties in experimental *in vitro* models, infecting mammalian LLC-MK2 cells and cells in the physiological HAE model. In this 3D model mimicking the human nasal mucosa, C-85473-derived HMPVs produced in DuckCelt®-T17 cells can infect and sustain viral propagation over time, demonstrating the conservation of the properties required to infect HAE cells. Based on these results, it appears that post-translation modifications and HMPV virion packaging provided by DuckCelt®-T17 avian cells are compatible with preservation of function and antigenicity of the HMPV F glycoprotein. Indeed, HMPVs produced in DuckCelt®-T17 cells show full infectivity in LLC-MK2 cells, a HAE model, and mice; this indicates the presence of a full functional F protein, which is crucial for viral attachment and entry into host cells ^20^. More importantly, as the HMPV F protein is known to be the major viral antigen ^38^, we validate here that our engineered Metavac® LAV candidate produced in the DuckCelt®-T17 cell line can induce a neutralizing antibody response against both homologous and heterologous HMPV strains, in accordance with the already demonstrated cross-protection potential of metapneumoviruses ^20^. Interestingly, murine neutralizing antibodies induced by HMPVs produced in LLC-MK2 cells also efficiently neutralized rC-85473-GFP viruses produced in DuckCelt®-T17 cells (Table 1), suggesting that there is a correct folding of the F protein at the virion surface.

From an industrial application perspective, we aimed to identify the best operating parameters that enable high-yield viral production while lowering costs and/or speed up the product harvest. We have shown that the MOI used should not be lower than 0.01 and the cell density should not be higher than 1×10^6^ cell/mL at the time of inoculation to achieve a maximum production yield within a time period compatible with cellular growth kinetics. We previously showed that a metabolic change occurs when the cell density is 3×10^6^ cells/mL or higher ^28^, which could explain loss of susceptibility to HMPV infection observed in DuckCelt®-T17 cells at higher cell densities. In contrast, infection at a lower cell density (0.5×10^6^ cells/mL) shows similar viral amplification kinetics to those seen with an infection at a cell density of 1×10^6^ cells/mL, which could be advantageous for upscaling the production process. Given these results, further explorations will be focused on cell metabolism, investigating whether different feeding strategies or a fed-batch approach could enhance or extend viral amplification.

As infection with C-85473 HMPV is known to be trypsin-dependent in adherent cell models because of the impact of trypsin on F protein properties ^32–34^, a particular effort was made to identify the optimal trypsin supplementation required to achieve a high yield in DuckCelt®-T17 cells. As anticipated, the presence of trypsin in the culture medium was critical for virus infection; HMPVs being basically unable to infect DuckCelt®-T17 cells in the absence of trypsin. Accordingly, adding trypsin twice during the process was sufficient to initiate and sustain the viral production, without adverse effects on DuckCelt®-T17 cell viability and cell growth.

In conclusion, the DuckCelt®-T17 cell line appears to be a promising platform for the manufacture of viral vaccines, and more particularly for our LAV candidate Metavac®, which is efficiently produced while maintaining its full replicative and immunizing properties in a mouse model. Considering the permissiveness of the DuckCelt®-T17 cell line to several influenza strains and vaccine seeds and its suitability for cultivation in a variety of suspension facilities, including single-use bioreactors up to 2 L of working volume ^28^, bioproduction processes based on the DuckCelt®-T17 cell platform would be scalable in order to reach large-scale virus propagation and cost-effective vaccine production in industrial volumes.

## Methods

### Cells and viruses

The DuckCelt®-T17 cell line (ECACC 0907703) was grown in suspension in OptiPRO™ Serum Free Medium (SFM, Gibco) supplemented with 1% penicillin/streptomycin (10,000U/ml, Gibco), 2% L-glutamin (Gibco), and 0.2% Pluronic F68 (Gibco), as previously described ^28^. The culture was performed at 37°C in a CO_2_ Kühner incubator (ISF1-X, Kühner) with 5% CO2 and 85% humidity. Agitation speed depended on the culture scale: 175 rpm for a working volume of 10 mL in TubeSpin® 50 mL (TPP^®^); 110 rpm from 20 to 500 mL of a working volume in Erlenmeyer shaker flasks (Erlenmeyer flask polycarbonate DuoCAP®, TriForest). Cells were passaged every 3 to 4 days at cell concentrations of 0.7×10^6^ cell/mL. LLC-MK2 cells (ATCC CCL-7) were maintained in minimal essential medium (MEM, Life Technologies) supplemented with 10% fetal bovine serum (Wisent) and 1% penicillin/streptomycin (10,000U/ml).

The wild-type (WT) A1/C-85473 strain of HMPV (GenBank accession number: KM408076.1) and two A1/C-85473-derived recombinant viruses were used in this study. The recombinant rC-85473-GFP (green fluorescent protein) virus, which is a GFP-expressing C-85473 WT counterpart virus, and the ΔSH-rC-85473-GFP virus (Metavac®), a recombinant virus from which the viral SH gene sequence is deleted, were generated by reverse genetics, as previously described ^20,33^. In order to constitute initial working viral stocks, both of these viruses were amplified onto LLC-MK2 monolayers in OptiMEM (Gibco) in the presence of 1% penicillin/streptomycin and acetylated trypsin (T6763, Sigma) and concentrated by ultracentrifugation, as previously described ^20,33^. Viral stocks were titrated onto LLC-MK2 cells at 50% tissue culture infectious doses (TCID_50_)/mL according to the Reed and Muench method ^39^.

### Infection and HMPV production in DuckCelt®-T17 cells

DuckCelt®-T17 cells in a working volume of 10mL in TubeSpin® 50 mL or 500 mL in 1 L Erlenmeyer shaker flasks were inoculated with HMPV in OptiPRO™ SFM (Gibco) supplemented with 1% penicillin/streptomycin (Gibco), 2% L-glutamin (Gibco), 0.2% Pluronic F68 (Gibco), and acetylated trypsin (T6763, Sigma). The viral production was monitored over a 10-day culture period by cell numeration, viability estimation, fluorescent microscopy (EVOS™ M5000 Cell Imaging System, Invitrogen), infectious TCID_50_ titer measurement ^39^, and infectivity quantification by flow cytometry ^20^. Briefly, 10 μl of the suspension was diluted in trypan blue and analyzed using a Countess™ II FL Automated Cell Counter. We then harvested and centrifuged a minimal sample of 1×10^6^ cells in suspension, supernatants were titrated as TCID_50_/mL and pelleted cells were fixed in a 2% formaldehyde solution to be analyzed by flow cytometry (FACS CantoII analyzer, Becton Dickinson). The percentages of infected GFP-positive cells in a minimum of 1×10^4^ total cells were measured with FACS Diva software.

To constitute concentrated DuckCelt®-T17-produced viral working stocks, the whole suspension of cells was harvested after 7–8 days of production, clarified by centrifugation at 2000 rpm, and then the supernatant was concentrated by ultracentrifugation as previously described ^20,33^. The pellet obtained was resuspended in OptiMEM and stored at −80°C for further use.

### Transmission electron microscopy

DuckCelt®-T17-produced HMPVs were harvested and concentrated by ultracentrifugation as previously described. Viral particles were then resuspended in NaCl (0.9%) and filtered at 0.45 μm. Suspensions were adsorbed on 200-mesh nickel grids coated with formvar-C for 10 min at room temperature. Then, grids with suspensions were colored with Uranyless (Delta Microscopies) for 1 min and observed on a transmission electron microscope (Jeol 1400 JEM, Tokyo, Japan) equipped with a Gatan camera (Orius 1000) and Digital Micrograph Software.

### *In vitro* replicative assay

Confluent monolayers of LLC-MK2 cells in 24-well plates were infected with rC-85473-GFP or Metavac® HMPVs produced in DuckCelt®-T17 cells in suspension or in adherent LLC-MK2 cells at a MOI of 0.01, as described previously ^33^. Supernatants of infected wells were harvested in triplicate every 24 h for seven days and endpoint TCID_50_/mL titrations were performed on each sample. After harvest, infected cell monolayers were fixed in 2% formaldehyde and colored with crystal violet solution.

### Infection of reconstituted human airway epithelium

*In vitro* 3D-reconstituted human airway epithelium (HAE), derived from primary nasal cells from healthy donors (MucilAir™), was purchased from Epithelix (Switzerland). Viral inoculum corresponding to a MOI of 0.1 was added onto ciliated cells and incubated for 2 h at 37°C and 5% CO_2_. Infections were monitored for the 12 days after viral inoculation (days post-infection, dpi); images of infected epithelium were captured by fluorescent microscopy at 5 dpi and apical washes with warm OptiMEM were performed at 2, 5, and 12 dpi in order to extract viral RNA (QIAamp® Viral RNA kit, Qiagen).

### Real Time-quantitative Polymerase Chain Reaction (RT-qPCR)

Amplification of the HMPV N gene from viral RNA samples was performed by quantitative RT-PCR using the One-Step SYBR™ GreenER™ EXPRESS kit (Invitrogen) and primers: N-Forward 5’-AGAGTCTCAATACACAATAAAAAGAGATGTAGG-3’ and N-Reverse 5’-CCTATCTCTGCAGCATATTTGTAATCAG-3’, as previously described ^20^. The calibration of HMPV N copies was assessed by amplification of a plasmid kindly provided by Dr Ab Osterhaus (Erasmus Medical Center, Rotterdam).

### Animal studies

Four- to six-week-old female BALB/c mice (Charles River Laboratories) were housed randomly in groups of five per microisolator cage. Twenty mice were infected by intranasal instillation with 1×10^6^ TCID_50_ of Metavac® viruses produced in LLC-MK2 adherent cells (Metavac® LLC) or in in-suspension DuckCelt®-T17 cells (Metavac® T17). As a control group, mice were mock-infected intranasally with OptiMEM. Animals (n=10) were monitored on a daily basis over 14 days for weight loss or presence of clinical signs. Mice were euthanized at 5 (n=4) or 14 dpi (n=6) using sodium pentobarbital and lungs were removed for the evaluation of viral titers. For virus titration, lungs were homogenized in 1 mL of phosphate-buffered saline (PBS) before N gene quantification by RT-qPCR.

To evaluate the induction of a neutralizing antibody response, mice were prime-infected intranasally with 5×10^5^ TCID_50_ of the rC-85473-GFP virus. Thirty days after prime infection, mice were boost-infected intranasally with 5×10^5^ TCID_50_ of Metavac® LLC or Metavac® T17 (n=10 per group). As control groups of immunization, a group of mice was prime-instilled with OptiMEM and boost-infected with 5×10^5^ TCID_50_ of rC-85473-GFP, and another group was prime-infected with 5×10^5^ TCID_50_ of rC-85473-GFP and boost-instilled with OptiMEM. Animals were monitored on a daily basis, and three mice were euthanized on day 5 after boost infection for the evaluation of viral titers in lung homogenates by RT-qPCR. To evaluate the production of neutralizing antibodies, blood samples were harvested by submandibular puncture prior to prime and boost infections (samples from five mice were pooled) and by intracardiac puncture 21 days after boost infection (n=6). Serial twofold dilutions of sera were then tested for neutralization of homologous rC-85473-GFP viruses produced in DuckCelt®-T17 cells (or in LLC-MK2 adherent cells, supplementary data) or neutralization of the heterologous WT CAN98-75 strain. Reciprocal neutralizing antibody titers were determined by an endpoint dilution assay, as previously described ^14^.

Animal studies were approved by the SFR Biosciences Ethics Committee (CECCAPP C015 Rhône-Alpes) under protocol number ENS_2017_019 and in accordance with the European ethical guidelines 2010/63/UE on animal experimentation.

### Statistical analysis

All statistical tests were conducted using GraphPad Prism5, comparing results expressed as the mean ± SD for each condition, using two-way ANOVAs with Bonferroni post-tests or one-way AVOVAs with Dunnett’s post-test.

### Data availability

All data generated or analysed during this study are included in this published article (and its supplementary information files).

## Supplementary materials

**Supplementary Table 1:** Induction of neutralizing antibodies against rC-85473-GFP (produced in LLC-MK2 cell line) by Metavac® viruses in mice

| Prime infection | Inoculum (log<sub>10</sub> TCID<sub>50</sub>) | Boost 30-day post-prime | Inoculum (log<sub>10</sub> TCID<sub>50</sub>) | Reciprocal neutralization titer (n=3)<sup>a</sup> |  |
| --- | --- | --- | --- | --- | --- |
|  |  |  |  | Against the rC-85473-GFP (LLC) virus |  |
|  |  |  |  | 29 days post-prime | 21 days post-boost |
| Mock | - | rC-85473-GFP | 5.7 | < 5 | 10 |
|  |  | Mock | - | < 5 | <5 |
| rC-85473-GFP | 5.7 | Metavac® LLC | 5.7 | 5 | >160 |
|  |  | Metavac® T17 | 5.7 | 5 | >160 |
|  |  | Mock | - | 5 | 5 |

<sup>a</sup>Three pools of sera from two mice were tested for neutralization against the rC-85473-GFP (LLC) virus, resulting in three biological replicates per group. One day before prime infection, the naïve status of mice was confirmed by a microneutralization assay from a pool of sera.

## Acknowledgments

We thank the microscopy service of the Centre d’Imagerie Quantitative Lyon-Est (CIQLE) in Lyon, the flow cytometry service of Plateforme de Cytométrie en flux at the Centre de Recherche en Cancérologie de Lyon (CRCL), and the animal care services of the Plateau de Biologie Expérimentale de la Souris in Lyon. We thank Fortune Bidossessi for her technical contribution to *in vivo* experiments (INRAE, UVSQ, VIM, 78350 Jouy-en-Josas, France) and Fanny Salasc for her technical contribution to cell culture and viral production (CIRI, Team VirPath U1111, Lyon, France).

## Author contributions

Conceptualization, J.D., G.B. and M.R.-C.; methodology, J.D., C.C., A.P. and M.R.-C.; validation, J.D., C.C. and A.P.; formal analysis, J.D., C.C. and A.P.; investigation, J.D., C.C., A.P., D.O.-M., A.T., P.B., B.P., E. L., C.M., V.D., T.J. and M.G.; resources, M.R.-C.; writing—original draft preparation, J.D., C.C. and M.R.-C.; writing—review and editing, A.P., O.T., J.F.-E., K.M., M.-È.H., M.R.-C. and G.B.; visualization, J.D., C.C.; supervision, B.L., G.B. and M.R.-C.; project administration, J.D. and M.R.-C; funding acquisition, G.B. and M.R.-C.

